# Distance-dependent consistency thresholds for generating group-representative structural brain networks

**DOI:** 10.1101/412346

**Authors:** Richard F. Betzel, Alessandra Griffa, Patric Hagmann, Bratislav Mišić

## Abstract

Large-scale structural brain networks encode white-matter connectivity patterns among distributed brain areas. These connection patterns are believed to support cognitive processes and, when compromised, can lead to neurocognitive deficits and maladaptive behavior. A powerful approach for studying the organizing principles of brain networks is to construct group-representative networks from multi-subject cohorts. Doing so amplifies signal to noise ratios and provides a clearer picture of brain network organization. Here, we show that current approaches for generating group-representative networks over-estimate the proportion of short-range connections present in a network and, as a result, fail to match subject-level networks along a wide range of network statistics. We present an alternative approach that preserves the connection-length distribution of individual subjects. Due to this simple modification, the networks generated using this novel approach successfully recapitulate subject-level properties, outperforming all existing approaches by better preserving features that promote integrative brain function rather than segregative. The method developed here holds promise for future studies investigating basic organizational principles and features of largescale structural brain networks.

## INTRODUCTION

The human brain is a network composed of neural elements – neurons, populations, and areas – interconnected to one another *via* synapses, axonal projections, and myelinated fiber tracts [1]. These connections shape neural elements’ patterns of input/output and play an important role in determining any given element’s functional properties [2]. By modeling neural elements and their connections as the nodes and edges of a graph, we can quantify with summary statistcs the network organization of brains and shed light on their function in health, disease, and development [3].

Though considerable effort has been expended to better understand how different aspects of brain network architecture vary across individuals [4] and covary with behavioral and clinical traits [5, 6], studying group-representative brain networks has also proven profitable for understanding the network organization and properties of a typical or average brain [7, 8]. Typically group-representative networks are generated by aggregating network data from many subjects while preserving those properties that are consistently expressed at the subject level [9–11]. This approach, when performed carefully, can theoretically enhance signal while suppressing noise and artifacts, affording a clearer view of the brain’s network organization.

Most methods for constructing group-representative networks are variants of “consistency-based thresholding.” That is, the group network is generated by specifying a threshold whose value ranges between 0 and 1, and retaining connections that are observed in at least a fraction of subjects. Next, each those connections are usually associated with a weight while all others are set to zero [9]. In almost every application, the same consistency threshold is applied uniformly over all possible connections. This so-called uniform consistency-based thresholding is common and group-representative networks generated using this approach appear frequently in the network neuroscience literature.

Group-representative networks are intended to serve as exemplars by preserving features consistently expressed at the level of individual subjects while reducing the level of noise and uncertainty. Among the most salient features of subject-level structural brain networks is the dependence of their topological features on their spatial embedding [12]. Both the probability that two brain areas are connected and the weight of that connection, should it exist, decay monotonically with inter-areal distance. This effect has been reported in human structural networks reconstructed from diffusion MRI with tractography algorithms [13–15], as well as networks reconstructed using invasive methods, e.g. tract-tracing [16, 17], suggesting that these dependencies are not simply artifacts of any specific network construction approach, but an evolutionarily conserved feature of large-scale brain networks [18].

The preference for strong, short-range connections can be explained parsimoniously by cost-reduction mechanisms. Intuitively, longer connections are more costly; they require additional material to form and extra energy for sustained use compared to short-range connections. As a consequence, nervous systems have evolved to favor shorter, low-cost connections. Despite this, brain networks still exhibit some long-distance connections. It is generally understood that longer connections play critical functional roles in order to offset their cost, though their precise function is still not fully understood.

Whatever their precise functional role, long-distance connections are arguably one of the most important subject-level features to preserve in a group-representative network. They play an important role in increasing shortest-path efficiency [19] and engender diverse network dynamics and information processing [14]. However, uniform consistency-based thresholding can produce networks that vastly under-estimate the number of observed long-distance connections. This bias emerges because the consistency of connections across subjects is, itself, distance-dependent, with short-range connections appearing more consistent than longer-range connections. As a consequence, for a given consistency threshold the distribution of supra-threshold connections will always favor short-range connections at the expense of long-distance connections. That is, group-representative networks generated using a uniform consistency-based thresholding procedure will exhibit more short-range connections and fewer long-distance connections than the typical subject (Fig. 1d-f). Because long-distance connections are responsible for driving certain network statistics, these group representative networks also fail to match subject-level networks in terms of those metrics.

**FIG. 1.**
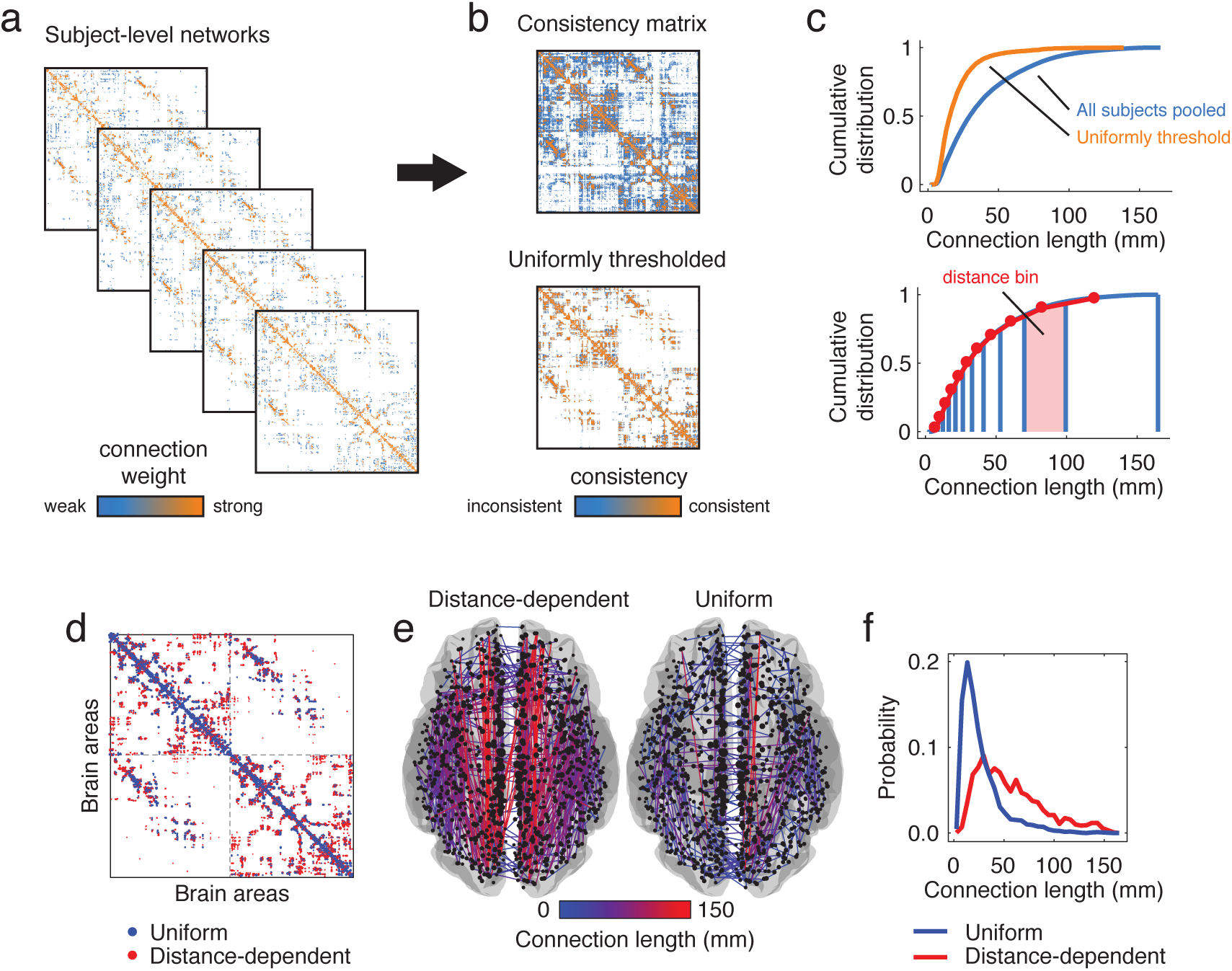
Construction and superficial comparison of group-representative matrices. Group-representative connectivity matrices summarize subject-level network data (*a*) by retaining features that are consistently expressed across subjects. In most applications the features of interest are the edges between brain areas and their weights. The most straightforward approach for generating a group-representative matrix involves first constructing a *consistency matrix* (*b*), whose elements denote the fraction of all subjects in which edges are expressed. Group-representative matrices can be estimated by retaining all connections expressed in at least *τ* subjects and populating those connections with weights. Though this approach is common, it suffers from a number of shortcomings. In general, because the probability of observing any given short-range connection is greater than the probability of observing a long-range connections, short-range connections also appear more consistently across subjects. As a result, imposing a *uniform consistency-based threshold* across all elements of the consistency matrix will result in a group-representative matrix in which short-range connections are expressed with much greater frequency than any single subject (*c*). To circumvent this issue, we present a simple alternative approach. Briefly, this involves dividing all connections into *m* bins according to their length and, within each bin, retaining the connection that is most frequently expressed. This *distance-dependent consistency-based thresholding* approach results in networks with almost the exact same edge length distribution as the typical subject. We also show differences in the group-representative matrices generated using the distance-dependent and uniform consistency-based threshold (here, we choose *τ* for the uniform model such the resulting matrix has a number of connections equal to that of the average subject) (*d*) Connections present only in the uniform model are depicted in blue and those present only in the distance-depdendent model are shown in red. (*e*) We plot these same model-specific connections on the brain, and color them according to their length (in millimeters). (*f*) In general, the connections unique to the uniform model are short-range (blue curve) while those unique to the distance-dependent model include long-distance connections.

Here, we present an alternative method for constructing group-representative networks. Our approach builds upon the consistency-based thresholding framework; rather than imposing a threshold uniformly over all connections, we allow our threshold to vary as a function of distance, retaining the most consistent connections conditional upon their length. We compare networks generated using this distance-dependent thresholding procedure with those generated using more traditional methods and show that, across a wide range of network statistics and comparative measures, networks generated using the distance-dependent approach outperform others. The distance-dependent procedure successfully recapitulates many of the important organizational features of subject-level networks and demonstrates promise for future exploratory studies of structural brain networks.

## RESULTS

In this section we compare four different approaches for generating group-representative structural connectivity networks.

- Connections are retained if they appear in at least one subject. We refer to this as the “Simple” model.
- Connections are retained if they appear in at least 50% of subjects. We refer to this as the “*τ* = 0.5” model.
- Connections are retained if they appear in at least *τ*_Avg_ subjects, where *τ*_Avg_ is the consistency threshold that results in a binary density equal to that of the typical subject. We refer to this as the “*τ* = Avg” model. Note: we calculate this threshold separately for inter-/intra-hemispheric connections.
- Connections are retained using a distance-dependent consistency threshold. The resulting network preserves, approximately, the edge length distribution of the typical subject. We refer to this as the “Dist.” model. As with the “*τ* = Avg” model, the distance-dependent threshold is introduced separately for inter-/intra-hemispheric connections.

This section is further divided into four subsections. In the first two subsections, we compare statistics of group-representative networks with those of individual subjects. In the next subsection, we characterize connectivity patterns of the group-representative matrices with respect to cognitive systems and discuss implications for our understanding of brain function. In the final subsection, we characterize how hubs are redistributed depending upon which of the four approaches for generating group-representative brain networks we choose.

Throughout this section, we report results of analyses using the high-resolution parcellation of the Lausanne dataset (*N* = 1000 nodes), where white matter fiber tracts are reconstructed from diffusion spectrum imaging data using deterministic streamline tractography (See *Materials and Methods* for processing details). These results are representative of our findings using coarser parcellations. Those additional results are included in the **Supplementary Materials**, Fig. S1, and Fig. S2.

### Uniform and distance-dependent consistency-based thresholding generate systematically different networks

The presence/weights of edges in structural connectivity networks exhibit spatial dependencies due to cost-reduction principles and reconstruction artifacts that cause short-range connections to be more consistently expressed across subjects. As a consequence, procedures for generating group-representative networks that retain connections using uniform consistency thresholds will necessarily over-estimate the number of short-range connections in a network.

In the following subsections we explore the implications of these biases in greater detail. Here, we simply show that uniform consistency thresholds generate group-representative networks with different spatial statistics than those generated using distance-dependent consistency thresholds, wherein the threshold for edge retention varies as a function of Euclidean distance. In Figure 1d, we show an adjacency matrix containing connections that are present in either the uniform or distance-dependent group-representative matrix but not both. Alongside this panel and in Figure 1e, we plot these same connections in anatomical space and color connections according to their lengths, with long/short connections appearing bright red/dark blue. In the left and right sub-panel we show connections that are present in distance-dependent model but not the uniform model and *vice versa*. Note that the connections shown in the left sub-panel tend to be long (red), indicating that the distance-dependent model retains long-distance connections that are not preserved in the uniform model. Conversely, the connections in the right sub-panel tend to be short (blue), indicating that the uniform model retains short-range connections not observed in distance-dependent model. These trends can be summarized by examining the distribution of connections present in one model but not the other. In Figure 1f we show that, as expected, connections retained exclusively by the uniform model are sharply peaked around 25 mm, whereas the connections retained by the distance-dependent model are more broadly distributed and include many longdistance connections.

These observations about differences in the connection length distributions of group-representative brain networks, though superficial, have important practical consequences for the structural properities of those networks. An over-expression of short-range connections could result in excessively cohesive brain network modules [13, 15], missing out on potentially rich intermodular connectivity patterns [20]. Conversely, group-representative networks that over-express long-distance connections may lack integratitve network properties such as local clustering [21]. These observations are in line with the fact that, in general, network properties are not independent of another, and variation in one property has implications for others [22]. Here, and throughout this paper, we argue that mischaracterizations of edge length distributions have profound implications for the spectrum of network properties that are exhibited by group-representative networks and whether those properties are in line with those of individual subjects’ networks.

### Consistency-based thresholding does not preserve subject-level network statistics

There are many criteria by which group-representative connectivity matrices could be evaluated and judged. Arguably among the most important is their ability to recover and recapitulate the topological properties of the subject-level data that they supposedly represent. In this section we compare four approaches for generating group-representative networks according to how well each matches individual subjects in terms of an ensemble of network statistics. We focus specifically on degree, strength, clustering coefficient, betweenness centrality, and edge length distribution and number of binary connections, total weight, mean clustering, topological efficiency, mean path length, modularity, diameter, and degree assortativity.

We divided network meausures into two categories based on whether they were defined locally or globally. We compared subject-level and group-representative local measures, i.e. those defined at the level of individual nodes, using Kolmogorov-Smirnov (KS) tests. The KS test statistic measures the maximum distance between two cumulative distributions and therefore smaller values indicate closer correspondence. In Figure. 2a-e, we overlay cumulative distributions of degree, strength, clustering coefficient, betweenness centrality, and edge length for each of the four group-representative on top of the cumulative distributions for individual subjects. The KS statistics comparing these distributions are plotted in Figure. 2f-j.

**FIG. 2.**
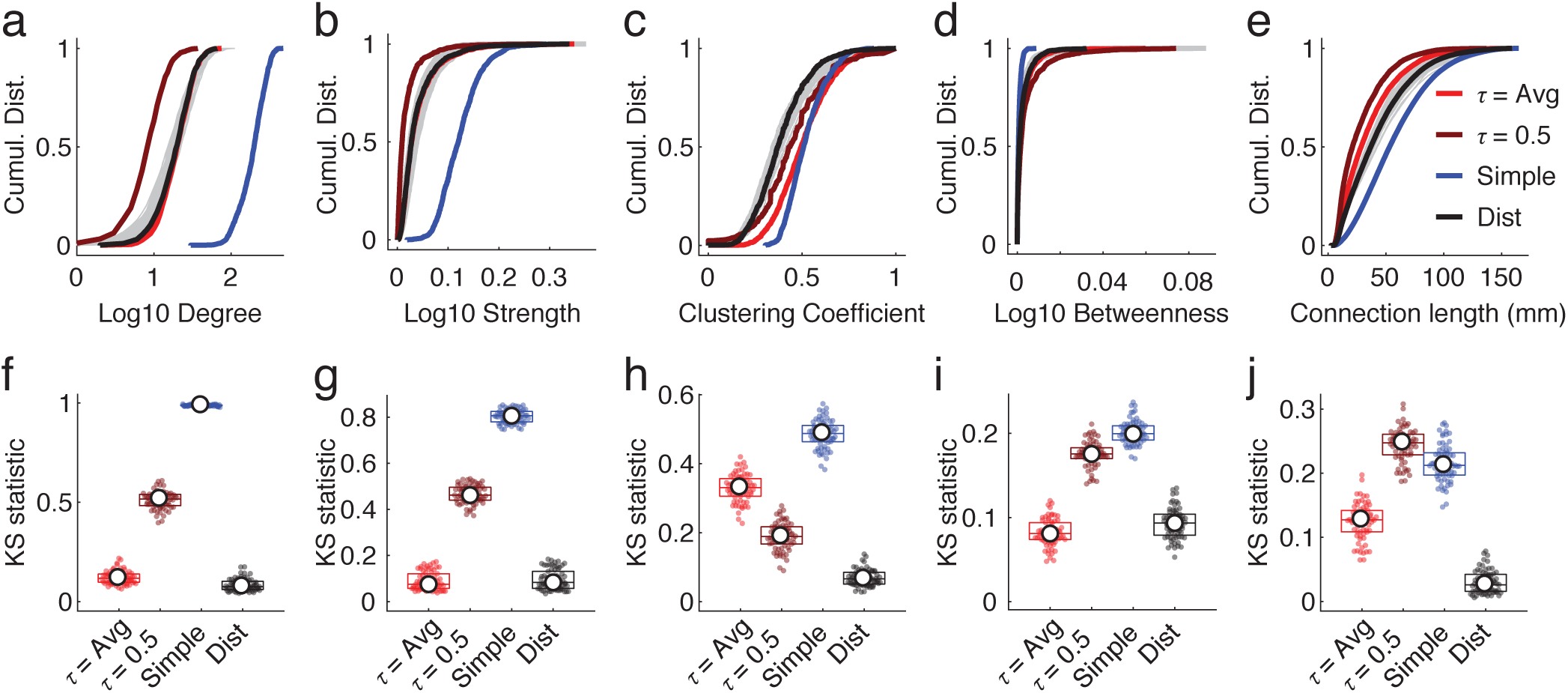
Comparing distributions across models. We show cumulative distributions for (*a*) degree, (*b*) strength, (*c*) clustering coefficient, (*d*) betweenness centrality, and (*e*) connection length. Subject-level data are shown in gray. Superimposed on those distributions are curves associated with the four models that we tested (in color). In panels (*f*) (*j*) we show KolmogrovSmirnov (KS) statistics for each network measure, which compare cumulative distribution curves of models with individual subjects.

For all five measures, we found that the distance-dependent consistency-based thresholding approach outperformed the other three models, i.e. smaller KS statistics (*p* < 0.05; Bonferroni corrected). These findings indicate that the distance-dependent model better preserved multiple node- and edge-level measures than the other models, suggesting that network statistics computed on the other group-representative networks may be misleading, in that they are not necessarily representative of the typical subject.

We performed similar comparisons of the global network measures. Here, rather than comparing distributions using the KS test, we z-scored the measures computed on the group-representative networks against the corresponding subject-level distributions. A z-score close to zero implied that the group-representative network was close to the mean subject-level value for a given network measure. In Figure. 3a-h, we show binary and weighted analogs of the total number of connections in the network, mean clustering, efficiency, and modularity. We found similar results when comparing network diameter, assortativity, and mean path length (Fig. 3i,j).

**FIG. 3.**
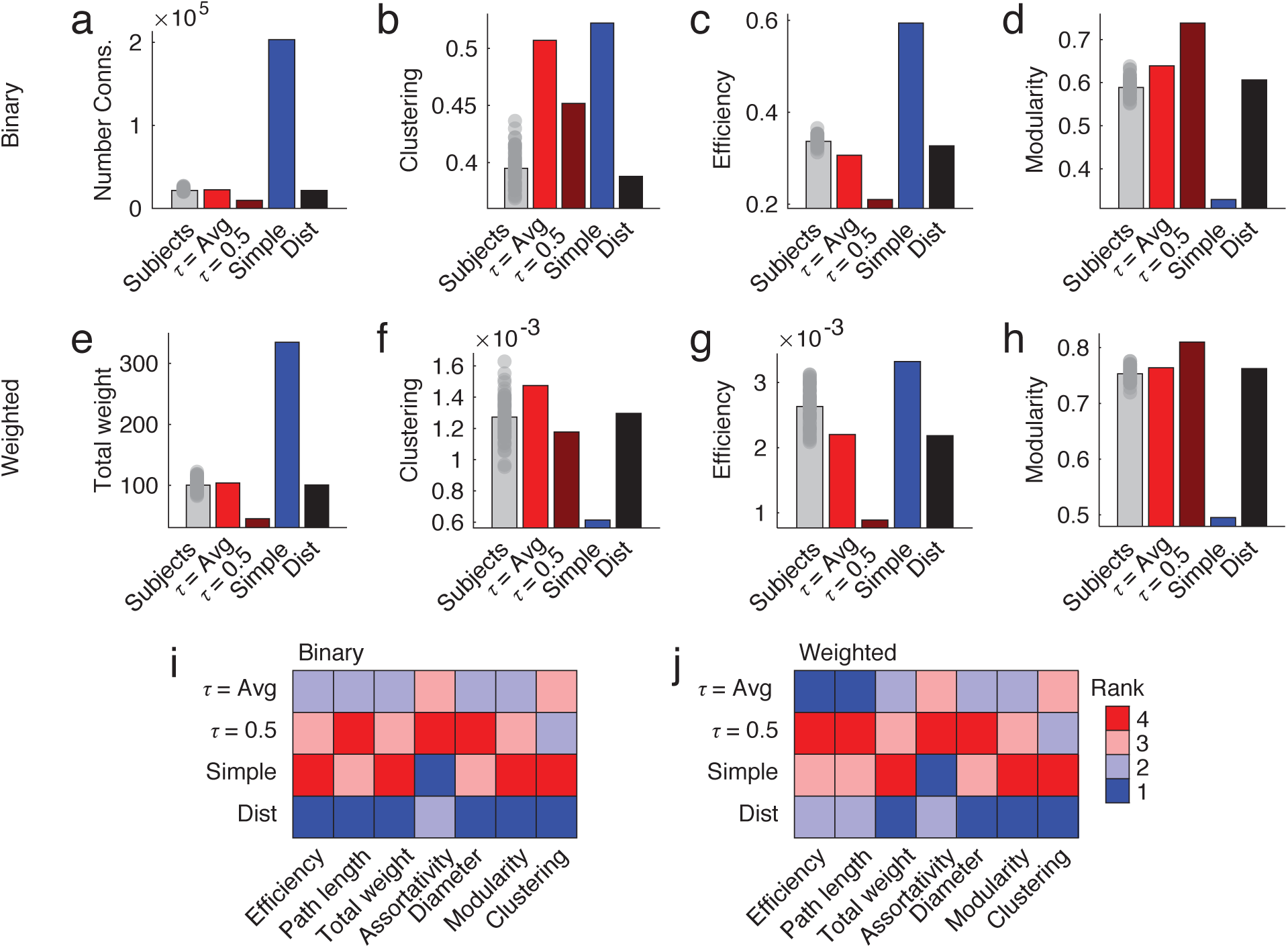
Comparing scalar network statistics. Here, we compare the performances of four different models of grouprepresentative brain networks to those of individual subjects. (*a*) Each bar represents the total number of binary connections for single subjects (gray), a uniform model with approximately the same number of connections as the average subject (bright red), a uniform model with a consistency threshold of *τ* = 0.5 (dark red), a “simple” model that retains a connection if its observed in even one subject (blue), and the distance-dependent model (black). Panels *b d* show similar plots but for mean clustering, efficiency, and modularity. Panels *e h* depict those same measures, but computed over weighted analogs of the binary networks. (*i*) For each measure shown (along with several others), we identified the model that was closest to that of the average across all subjects. In general, we find that the distance-dependent model consistently outperforms or performs comparably to the other tested models, achieving rank 1 or 2 across all metrics.

As with the local network measures, these findings suggest that decisions about how to generate a group-representative connectivity matrix have implications for its topological organization. Importantly, the most popular approach – uniform consistency-based thresholding preserves a greater number of short-range connections compared to the typical subject and, as a result, exhibits topological properties that are inconsistent with those exhibited by that subject.

### Implications for structure-function relationships

Another dimension along which group-representative structural networks can be compared is the extent to which they match subject-level networks in terms of their structure-function association. Here, we map brain structure to function by averaging the strengths of connections within and between previously described cognitive systems – resting-state networks (RSNs) [23]. Restingstate networks or (intrinsic connectivity networks) are composed of brain areas with similar *functional* connectivity profiles and maintain a close correspondence with patterns of areal activation under different cognitive task. We do this for each of the four group-representative models, generating a system-by-system connection density matrix (Figure. 4b). We also do the same for each individual subject, which we then average to obtain a single density matrix describing inter-system connection densities of subjects (Figure. 4a). We compare group and subject matrices as the correlation of their elements to one another.

**FIG. 4.**
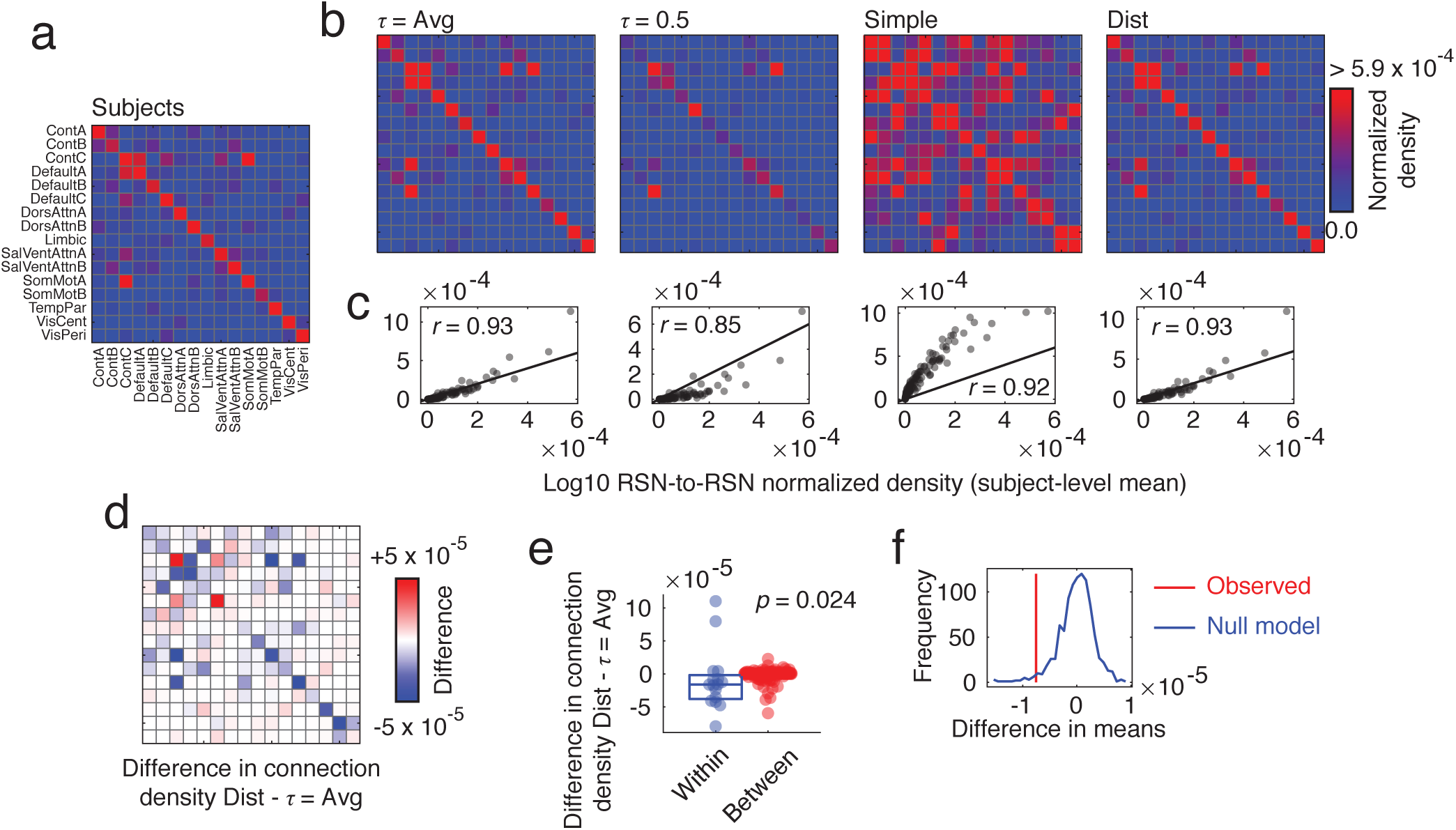
Comparing within-/between-RSN connectivity patterns. We compared different group-representative networks in terms of connection densities within and between canonical brain systems taken from [23]. (*a*) Inter-RSN connection density of the typical subject. (*b*) Inter-RSN connection densities for four different group representative networks: (from *left* to *right*) uniform consistency model with same density as subjects, uniform consistency model with threshold set at *τ* = 0.5, “simple” model, and distance-dependent model. (*c*) We show the correlation patterns of inter-RSN densities for each model (*y* axis) with that of the subject average (*x*-axis). Of the models compared here, the distance-dependent and the uniform model with same density as the typical subject performed the best. We compare these models so as to better understand their differences. (*d*) Difference in inter-RSN connection density between. Blue colors indicate that connection density is greater in uniform model while red density indicates that connection density is greater in distance-dependent model. (*e*) We find that, on average, the uniform model results in weaker within-RSN density than the distance-dependent model, while the distance-dependent model has greater between-RSN density. (*f*) We show the observed difference in withinand between-RSN density and compare it against a null model. Here, we show the null distribution (blue) and the observed value (red). The null distribution was constructed by independently and randomly permuting rows/columns of each original connectivity matrix and re-aggregating according to the RSN system labels. Then we compute the mean difference of within-/between-RSN densities.

In general, we find that each group-representative matrix is positively correlated with the subject-level matrix, indicating that, overall, system-to-system connectivity at the subject level is preserved at the group level by all models. Nonetheless, there is considerable variability across group-representative models in terms of correlation magnitude and deviation from the identity line (Figure. 4c). For instance, the “simple” model exhibited a correlation of *r* ≈ 0.92 but massively over-estimated the amplitude of connection densities. Similarly, the uniform model with a threshold of *τ* = 0.5 exhibited a much weaker correlation of *r* = 0.85. Whereas, the other models exhibited much stronger correlations with magnitudes in excess of *r* ≈ 0.93.

The two best-performing models were the uniform model using a threshold that resulted in the same density as the typical subject and the distance-dependent model, which we compared in greater detail. First, we computed the difference in inter-RSN connectivity density (Figure. 4d). We found that there were subtle yet systematic differences between the two models. In particular, we found that the distance-dependent model exhibited much weaker within-RSN density compared to that of the uniform model while also exhibiting stronger between-RSN connection density (*p* < 0.05, permutation test; Figure. 4e, f).

These findings have important implications for the analysis and interpretation of brain network data. This is especially true for studies that aim to link features of structural and functional brain networks to one another. Past studies using group-representative data constructed using a uniform consistency threshold fail to match the specificity of subject-level networks while the simple averaging procedure over-estimates the weights of connections. These failings can lead to mischaracterizations of structure-function associations.

More importantly, these findings suggest that differences in the construction method for group-representative networks can result in networks that emphasize either segregative features – i.e. stronger within-RSN connection densities, as expressed by the uniform model – or integrative features – i.e. stronger between-RSN connection densities as expressed by the distance-dependent model. The balance between information segregation and integration is thought to be an important organizational principle responsible for shaping brain network topology [24–26]. Our findings indicate that different group-representative models differentially emphasize these characteristics, indicating that a user’s seemingly arbitrary choice in model can have implications for measures made on a network.

### Hub (re)distribution

A third means of comparing group-representative networks against one another is to measure the redistribution of hub areas, i.e. assessing changes in the locations of “central” brain areas as a result of choosing one group model *versus* another. Here, we compare the spatial distribution of betweenness centrality, node degree, clustering coefficient, and participation coefficient under uniform and distance-dependent models. To ensure that comparisons are as fair as possible, we rank-transformed all measures prior to comparison.

In general, we found widespread and hemispherically symmetric re-distribution of hub regions. In the case of betweenness centrality (Figure. 5a), we found that under the distance-dependent model, areas associated with cognitive control are increasingly central, while areas in the somatomotor system become less central (*p* < 0.05; corrected for multiple comparisons by controlling FDR at 5%; Figure. 5e). In terms of degree (Figure. 5b), we find that control and limbic systems make a greater number of connections, while dorsal attention, salience/ventral attention, somatomotor, and visual systems exhibit fewer connections (Figure. 5f). For clustering coefficient (Figure. 5c), we find that components of default mode and motor systems are more clustered while multiple components of the control network are less clustered (Figure. 5g). Finally, in terms of participation coefficient (Figure. 5d), we find that the dorsal attention and visual systems exhibit greater participation whereas somatomotor and other visual systems exhibit decreased participation (Figure. 5h).

**FIG. 5.**
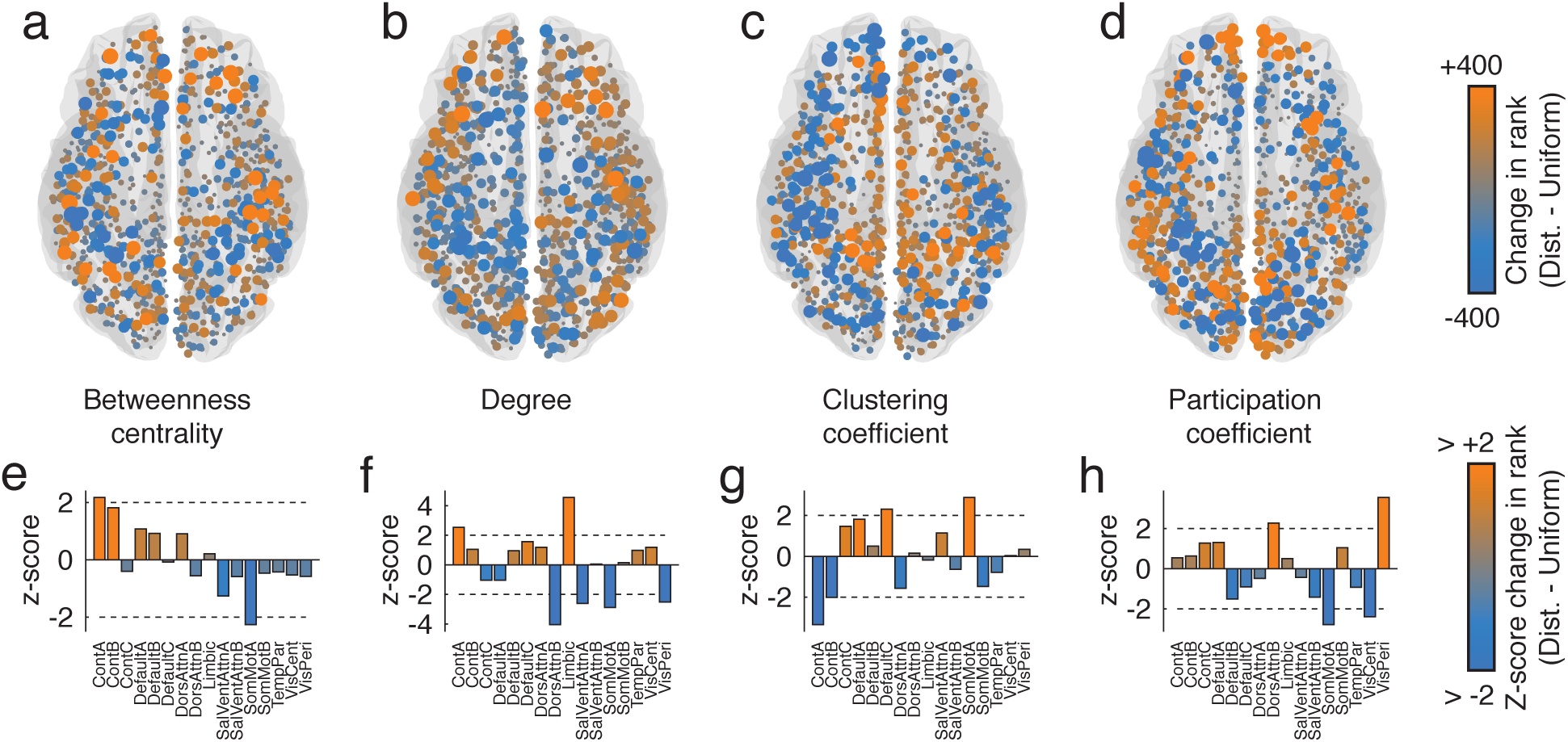
Comparing spatial distribution of hubs. We compare four measures of hubness –(*a*) betweenness centrality, (*b*) degree, (*c*) clustering coefficient, and (*d*) participation coefficient. Rather than compare raw values, which can fluctuate due to small differences in global network properties like total number of connections or weight, we compare ranked values of each measure and observe whether a node’s rank is smaller/greater under the distance-dependent or uniform model. Orange-colored nodes indicate that a node’s value is greater under the distance-dependent model than it is under the uniform model. Bluecolored nodes indicate the opposite. We then aggregated node-level differences in ranked measures by cognitive systems and compare the mean system-level values with those obtained under a null model. In panels *e*-*h* we show the z-scored system means. In general, large-magnitude z-scores indicate bigger greater system-level differences between the two models.

These findings have important implications for our understanding of brain function. Hubs and central brain areas are believed to be important for controlling interareal communciation and regulating the flow of information within and between brain network modules. Indeed, the designation of an area as a “hub” has been important for hypothesis generation and has also played an important confirmatory role in other studies. Our findings suggest that these definitions are, at least to some extent, dependent upon the method used to generate a group-representative network. Moreover, some of the most salient differences between methods appear localized to specific cognitive systems, which has additional implications for how we interpret findings related to hubs and brain function.

## DISCUSSION

In this paper we develop a novel method for generating group-representative brain networks that outperforms more conventional methods in terms of preserving subject-level statistics. Specifically, we show that the new method better preserves local and global network statistics, that its structure-function relationships are more consistent with those of individual subjects, and that it gives rise to a different intuition of where highly central hub regions are located in the brain.

### Structural networks need long-distance connections

Here, we found that compared to uniform consistency-based thresholding, a distance-dependent threshold preserves to a greater degree the connection length distributions observed in individual subjects. Specifically, the uniform model retains a greater proportion of short-range connections than the typical subject. We argue that this difference in connection length distributions has both practical (i.e. measurable) and theoretical consequences. Practically, we show that expressing fewer long distance connections results in networks that are more clustered and, as a result of increased rate of triadic closure, more modular than that of the typical subject [27]. Similarly, lacking long-distance shortcuts results in networks that are less efficient and that possess longer characteristic path length than the average subject [13, 27]. Overall, the uniform model results in networks that emphasize segregative traits at the expense of those that support integration of information [24]. This is confirmed further when we compared the inter-system connection densities of the distance and uniform models, observing that within-community density was less on average, using the distance model compared to the uniform model. Additionally, the differences in the features preserved by each model contribute to shaping the spatial distribution of hubs across the brain. Overall, these findings suggest that the principle advantage of the distance-dependent model is that it better preserves network features that emphasize information integration.

### The role of group-representative network analysis

In this study we focus on group-representative networks. Analysis of these group networks has been and remains and important component of network neuroscience. In the case of non-human datasets, group network analysis is almost always performed out of necessity. Invasive methods like tract-tracing limit the number of experiments that can be performed on any one animal brain. As a result, whole-brain networks are necessarily composites of many animals [28–30]. Human structural networks constructed from diffusion MRI data using tractography algorithms are sensitive to scan parameters and prone to false positives and negatives [31,32]. Analyses of human networks, therefore, benefit from aggregation of multi-subject cohorts into a group-representative network, which serves to enhance signal while reducing the level of noise and uncertainty. The resulting networks can be treated as exemplars and used to uncover key structural traits and organizing principles [7,8], as the basis for dynamic models [33], and serve as a sort of “prior” for other machine and statistical learning approaches [34].

However, analysis and interpretation of group-level networks presume that those networks are, in fact, representative of the typical subject. Group networks that violate this assumption can contribute misleading or inaccurate insight into brain network organization and function. We show here that group-representative networks constructed using a uniform consistency-based threshold, which fail to preserve important spatial properties of subject-level brain networks, may be especially susceptible to such inaccuracies. Our work suggests that the uniform model generates networks that tend to overestimate the cohesiveness of communities. In addition, the uniform model also presents a conflicting account of hub distributions throughout the brain when compared with the distance-dependent model. Because analysis of group-representative networks remains a powerful approach, understanding and accounting for their limitations and biases should be investigated in future research.

### Limitations

Here, we present a novel method for constructing group-representative networks, demonstrating that this approach results in group networks that better preserve subject-level properties than existing approaches. Nonetheless, our study suffers from some limitations.

First, we make the overarching assumption that the long-distance connections observed in single-subject networks (which we preserve in our group-representative network) are “real” and not strictly artifacts of the tractography algorithm. This assumption is supported, first, by the fact that long-distance connections are, in general, more challenging to reconstruct using common tractography parameters. Completing long streamlines require strong spatial coherence of the diffusion field over distances greater than 150 mm, which is unlikely to occur in the presence of high background noise [35]. Second, long-distance connections, typically, do not appear randomly distributed, but are clustered [14]. That is, if regions *i* and *j* are connected by a long-distance tract, it is likely that other regions in *i*’s spatial neighborhood are connected to *j* and *j*’s spatially proximal neighbors (and *vice versa*). These observations suggest that longdistance connections cannot easily be explained as errant “one-off” reconstructions. Nonetheless, tractography has known shortcomings [31,32], and the verisimilitude surrounding long connections remains unclear. Advances in hardware, fiber reconstruction software [36], and detailed comparisons of tractography with tract-tracing data [37] will help future studies overcome these issues.

A second limitation concerns the network measures and metrics used to compare group-representative networks to one another and to individual subjects. These measures were selected because they emphasized network topology as well as its relationship to neuroscientifically relevant meta-data (i.e. cognitive functional systems). However, these measures are, first, not necessarily an exhaustive list and it is unclear whether the distance-dependent model’s performance would be better than other models were we to select a different set of measures. Second, network measures tend to be correlated with one another – e.g. a network with high efficiency will tend to have short-path length. Therefore, the comparisons we made were not necessarily independent of one another. Though we intentionally selected a wide range of measures to help address these issues, our analyses could be extended in future work to include a broader range of measures and comparative metrics.

A final limitation is the necessity that the subject-level matrices used to estimate the group-representative network be sparse. Both the uniform and distance-dependent models rely on the intuition that some connections are more common across individuals than others. For some diffusion MRI and tractography algorithms e.g. probabilistic tractography – this is not always the case [38]. Nonetheless, it may be possible to adapt the approaches used here with sparse deterministic tractography to the probabilistic case by substituting connection probability measures for the consistency. Care would have to be taken to deal with the potentially confounding geometric and spatial biases [15]. Future work should investigate this in greater detail.

## Conclusion

Overall, our findings suggest that care must be taken when studying and analyzing group-representative networks. We presented an approach for limiting discrepancies between subject- and group-level networks by adding a distance-dependence to the consistency threshold. This approach will aid in future studies that seek to investigate general properties of structural brain networks.

## MATERIALS AND METHODS

### Connectome dataset

In this study we compared methods constructing group-representative brain networks from structural connectivity data. We carried out these comparisons using diffusion spectrum MRI data parcellated into networks at three different organizational scales. Here, we describe those processing steps in greater detail.

#### MRI acquistion

70 healthy participants (age 28.8 ±9.1yo, 43 males) were scanned on a 3T scanner with a 32-channel head coil (Magnetom TrioTim, Magnetom Prisma, Siemens Medical, Germany). The session included (1) a magnetization-prepared rapid acquisition gradient echo (MPRAGE) sequence (1 × 1 × 1.2 mm resolution, 240 × 257 × 160 voxels; TR = 2300 ms, TE = 2.98 ms, TI = 900 ms); (2) a diffusion spectrum imaging (DSI) sequence (2. ×22×.23 mm resolution; 96 ×96 ×34 voxels; TR = 6100 ms, TE = 144 ms; q4half acquisition with maximum b-value 8000 s/mm^2^, one b0 volume). Informed written consent was in accordance with institutional guidelines and the protocol was approved by the Ethics Committee of Clinical Research of the Faculty of Biology and Medicine, University of Lausanne, Switzerland.

#### MRI preprocessing

MPRAGE volumes were segmented into white matter, grey matter and cerebrospinal fluid using FreeSurfer software version 5.0.0 [39]. Cortical volumes were segmented into five progressively finer parcellations, with 68, 114, 219, 448 and 1000 approximately equally-sized parcels [40]. Here, we analyze the 68-, 219-, 1000-parcel divisions. DSI data were reconstructed following the protocol described by Wedeen and colleagues [41], thus estimating an orientation distribution function (ODF) in each voxel. Up to three main streamline orientations were idenntified in each voxel as the maxima of the ODF (DiffusionToolkit software, http://www.trackvis.org/dtk).

Structural connectivity matrices were estimated for individual participants using deterministic streamline tractography on reconstructed DSI data, initiating 32 streamline propagations per diffusion direction per white matter voxel [42]. The MPRAGE and the brain parcellation were linearly registered to the subject diffusion space (b0) using a boundary-based cost function (FreeSurfer software) [43]. For each starting point, streamlines were grown in two opposite directions with a fixed step size equal to 1 mm. As the streamline entered new voxels, growth contributed along the ODF maximum direction that produced the least curvature. Streamlines were terminated if changes in direction were greater than 60 degrees/mm. Tractography completed when both ends of the streamline left the white matter mask. Structural connectivity between pairs of parcels was estimated in terms of streamline density, defined as the number of streamlines between two parcels normalized by the mean length of the streamlines and the mean surface area of the parcels.

### Single-subject networks and connection consistency

Let **A**_*s*_ ∈ ℝ ^*N* ×*N*^ be the weighted and symmetric structural connectivity matrix for subject *s* = 1, *…, T*, whose element *A*_*ijs*_ indicates the normalized streamline count between brain areas *i* and *j*. Given the set of matrices = {**A_*S*_**}we can calculate the consistency matrix,**C** ∈ ℝ ^*N*×*N*^, whose element 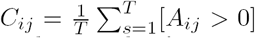. In tuitively, then, *C*_*ij*_ ∈ [0, 1] indicates the fraction of *T* subjects for whom the connection {*i, j*} is expressed.

### Group-representative network construction

In this paper we compare several strategies for constructing group-representative networks for structural connectivity estimated from dMRI and reconstructed using tractography. In this section, we introduce several approaches for doing so.

#### Simple average

The most näive approach for generating a group-representative connectivity matrix is to let each connection’s weight be its mean value over all subjects, ignoring those for whom a connection is not expressed, i.e. *A*_*ijs*_ = 0. We refer to this approach as the *simple average* and denote the estimated group-representative connectivity matrix as **A**^*simp*^, whose elements are defined as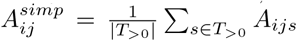Here, *T*_>0_ is the set of subjects satisfying *A*_*ijs*_ > 0.

#### Consistency-based thresholding

A more common approach for generating group-representative matrices is to impose a threshold, *τ*, over the elements of a consistency matrix, **C**, so that the only elements retained are those that satisfy *C*_*ij*_ > *τ*. The intuition is that a good group-representative matrix should preserve the features, in this case connections, that are consistently expressed across individual subjects. Within the broader consistency-based thresholding framework there are many strategies for implementation. In this section we hightlight those that are explored in this paper.

The most common variant of consistency-based thresholding is the imposition of a *uniform* threshold over all connections. That is, all possible elements of the consistency matrix are considered simultaneously and those that survive *C*_*ij*_ > *τ* are retained. In contrast, a *restricted* threshold is one in which connections are grouped into *K* classes according to some criteria and a class-dependent threshold, *τ* (*k*), where *k* ∈{1, *…, K}*, is imposed separately over each class. For example, connections could be classified according to whether their starting and termination points fall within the same or different hemispheres. The restricted threshold is not limited to ordinal data, but can also be used with continuous variables through discretization. Inter-areal distance, for instance, is a continuous variable that measures the Euclidean distance between areal centroids. One could impose a distance-dependent threshold, *τ* (*D*), by discretizing the interval of possible Euclidean distances into nonoverlapping bins. Each bin would include connections that span a particular range of distances, and distinct thresholds could be imposed within each bin.

#### Models tested in this submission

Here, we tested four different methods for generating group-representative networks. The first was the “simple” model, which retained a connection and its average weight if it was observed in at least one subject. The second and third models were variants of the uniform consistency-based threshold model. The first of these imposed a threshold of *τ* = 0.5 over all connections, so that the group network included only those connections expressed in at least half of the subject cohort. The second model involved choosing a consistency threshold such that resulting network had a binary density as close as possible to that of the average subject. We refer to this model as the *τ* = Avg model. The fourth and final model was a distance-dependent threshold, imposed exactly as described above.

### Network measures

We compared subject-level and group-representative networks using a set of measures that quantify different aspects of network topology (all measures computed using functions provided as part of the Brain Connectivity Toolbox https://sites.google.com/site/bctnet/; [44]). These measures included: binary and weighted total connection weight, degree, clustering coefficient (nodal and global), betweenness centrality, efficiency, path length, diameter, multi-scale modularity, participation coefficient, assortativity, and connection length. In this section, we describe those measures in greater detail.

#### Degree and total weight

Among the simplest structural measures one can calculate given a connectivity matrix, **A** = {*A*_*ij*_ *}*, is the degree of node *i*, which summarizes the total number (or weight) of its connections:

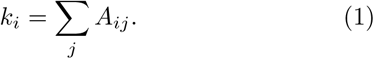

Given the vector of nodes’ degrees, **k** = {*k*_*i*_ }, we can then calculate the total weight of the network as 2*m* = Ʃ*i k*_*i*_ (the factor of two is necessary in this case due to the undirectedness of the networks considered here).

#### Clustering coefficient

Another simple measure is clustering coefficient of each node, *i*. The clustering coefficient measures the density of connections among all of *i*’s neighbors and is calculated as:

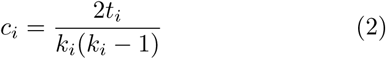

Where *t*_*i*_=Ʃ _*j,h*_*A*_*ij*_*A*_*ih*_*A*_*jh*_ is the number of triangles surrounding node *i*. The node-level clustering coefficient can be averaged to summarize the mean clustering of a network, 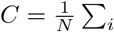

#### Path based measures: Characteristic path length, diameter, betweenness centrality, and efficiency

Given the connectivity matrix **A** = {*A*_*ij*_ }, we define the matrix **D** = {*D*_*ij*_ }to be the shortest paths matrix, whose element *D*_*ij*_ is equal to the length of the shortest topological path between nodes *i* and *j*. For binary networks, shortest paths are calculated in terms of geodesic distance, i.e. number of steps. For weighted networks, however, shortest paths are calculated based on a transformation of edge weights to length. Here, we use a reciprocol weight-to-length transformation, i.e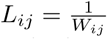Once the shortest paths matrix has been calcualted, we can define characteristic path length as:

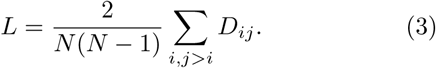

We can also define network diameter as:

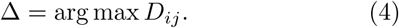

Finally, we define efficiency as:

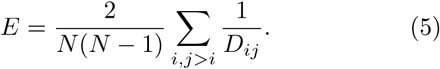

A measure related to the concept of shortest paths is betweenness centrality. Let *ρ*_*hj*_ be the number of shortest paths between nodes *h* and *j* and *ρ*_*hj*_(*i*) be the number of shortest paths between *h* and *j* that pass through node *i* Then the betweenness centrality of node *i*, which measures the fraction of all shortest paths that pass through node *i*, is calculated as:

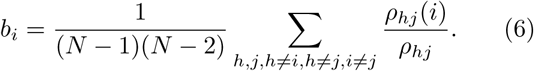

#### Modularity maximization and the participation coefficient

Many real-world networks exhibit modular architecture, meaning that their node’s can be meaningfully decomposed sub-networks (also called “communities” or “modules”) that are internally cohesive but segregated from one another. Though there are many approaches for detecting modules in networks, one of the most popular is modularity maximization, which partitions each node *i* into one of *K* communities σ_*i*_ ∈{1, *… K*} by maximizing a modularity quality function, designed by the variable *Q* [45]. Modularity functions have the following form:

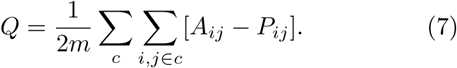

Here, *A*_*ij*_ and *P*_*ij*_ are the observed and expected weights of the connection between nodes {*i, j}*. The double summation ensures that, effectively, the sum counts only pairs of nodes that fall within communities, i.e. σ_*i*_ = σ_*j*_. Accordingly, *Q* is optimized when the observed density of connections within communities is maximally greater than what would be expected by chance. Here, we let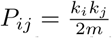, which corresponds to a chance model in which each node’s degree is preserved but where its connections are otherwise made at random.

The standard formulation of modularity, *Q*, suffers from what is known as a “resolution limit” rendering it incapable of detecting communities below some characteristic scale determined by a network’s overall density [46]. To circumvent this issue, a parameterized version of the modularity function exists:

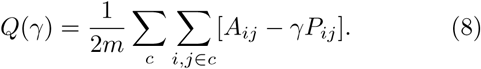

Here, γ is the structural resolution parameter, that scales the relative contribution of the expected weight. Effectively, the value of γ determines the scale of detected communities, with small and large values of γ returning correspondingly larger or smaller communities. This parameterization does not directly address the issue of the resolution limit; it simply shifts the scale below which communities are undetectable.

Optimizing *Q*(γ) is computationally intractable. However, there exist many methods and heuristics for approximating the optimal solution. One of the most common is the so-called “Louvain algorithm,” which has proven both fast and accurate in benchmarking tests [47]. The “Louvain algorithm” is stochastic, however, and its estimate of the optimal partitions depends upon initial conditions. Accordingly, it is common to repeat the algorithm many times to generate a sample of near-optimal solutions.

Here, we vary γ over a range of 0.7 to 2.1 in increments of 0.1. At each discrete value, we optimize *Q*(γ) using a generalization of the Louvain algorithm 100 times [48].

The partitions generated by modularity maximization and related community detection algorithms can be used to further characterize different aspects of network organization and function. One such metric is the participation coefficient, which measures the extent to which a node’s connections are distributed across modules or concentrated within its own module [49]. Let *?*_*iσ*_ denote the total weight of connections node *i* makes to module σ. Participation coefficient is calculated as:

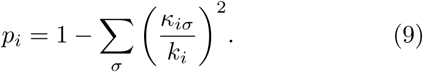

Intuitively, the closer *p*_*i*_ is to 1, the greater the extent that *i*’s connections are distributed across many different modules. Values of *p*_*i*_ close to 0 indicate that *i*’s connections are conncentrated within a few modules.

#### Degree assortativity

We also computed degree assortativity, a measure that quantifies the extent to which nodes of a given degree tend to connect with other nodes of similar degree [50]. Intuitively, degree assortativity is a Pearson correlation of the degrees at different endpoints of each edge in the network and is calculated as:

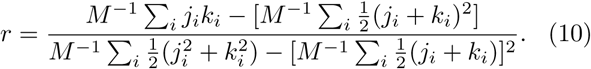

Here, *i* = 1, *…, M*, indexes the edges in the network, and *j*_*i*_ and *k*_*i*_ are the degrees of nodes connected by the *i*^th^ edge.

#### Connection length

Finally, we also computed connection length distributions. We define the length of a connection between nodes *{i, j}* as the Euclidean distance separating the centroids of those regions. In general, connections are curvilinear and do not adhere to straight-line distances. However, in studies that measured both Euclidean distance and the curvilinear fiber lengths of connections, these measures were found to be highly correlated [51]. This implies that, while Euclidean distance is not a perfect substitute for a connection’s length, it is a very good first-order approximation.

## CODE AVAILABILITY

Code for generating the uniform and distance-dependent group-representative networks is available at https://www.richardfbetzel.com/code/.

## ACKNOWLEDGMENTS

This research was undertaken thanks in part to funding from the Canada First Research Excellence Fund, awarded to McGill University for the Healthy Brains for Healthy Lives initiative. BM acknowledges support from the Natural Sciences and Engineering Research Council of Canada (NSERC Discovery Grant RGPIN #017-04265), the Fonds de recherche Québec Santé (Chercheur Boursier) and the Canadian Institutes of Health Research (CIHR; Project Grant #391300).

---

**FIG. S1.**
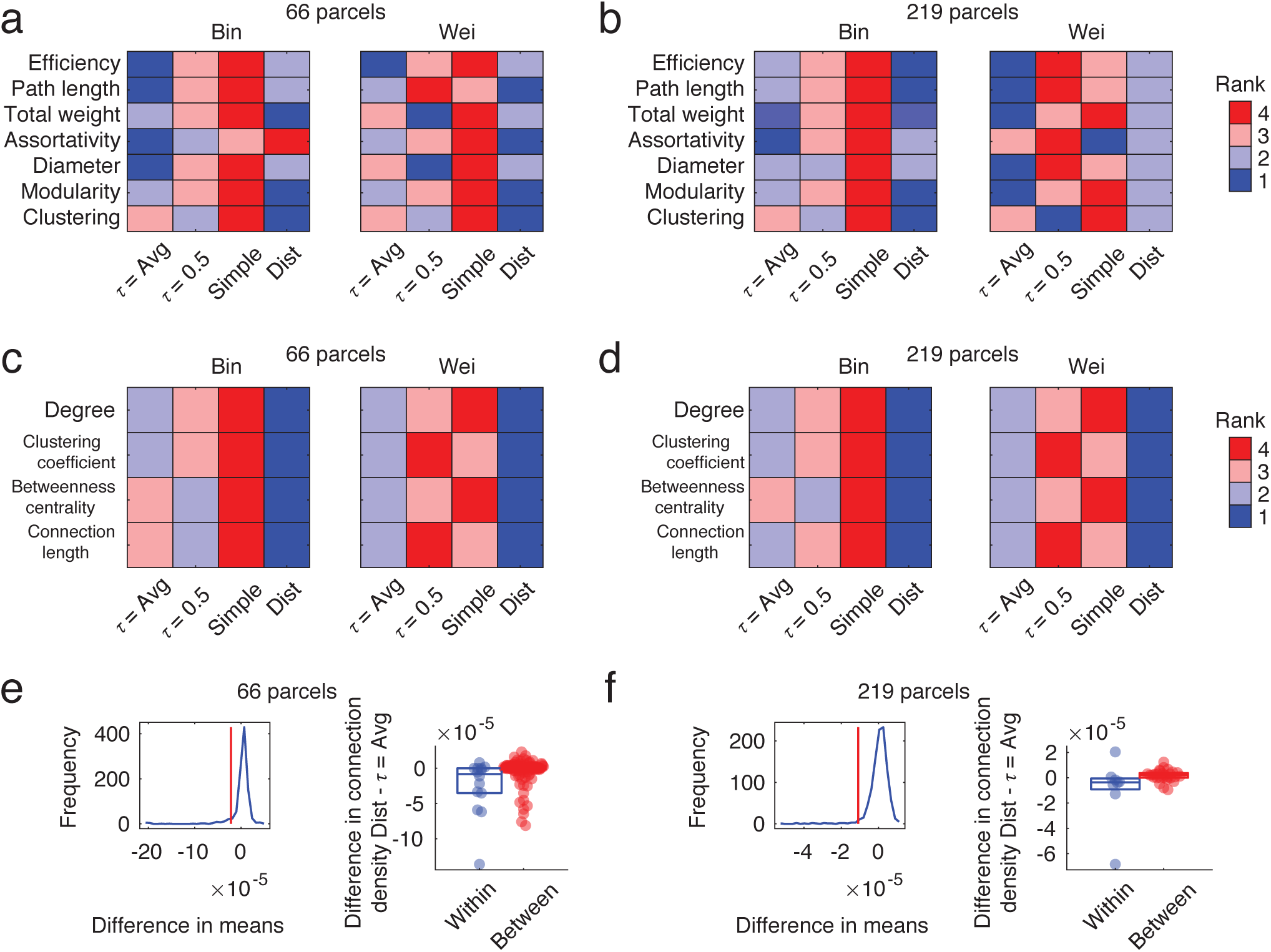
Similar results with different parcellation resolutions. In the main text we analyze networks of *N* = 1000 nodes. Here, we recapitulate those analyses using networks of *N* = 66 and *N* = 219 nodes. In panels *a* and *b*, we show results analogous to those shown in Fig. 3. In panels *c* and *d*, we show results analogous to those shown in Fig. 2. Finally, in panels *e* and *f*, we show results analogous to those shown in Fig. 4. In general, these supplementary results support those presented in the main text.

**FIG. S2.**
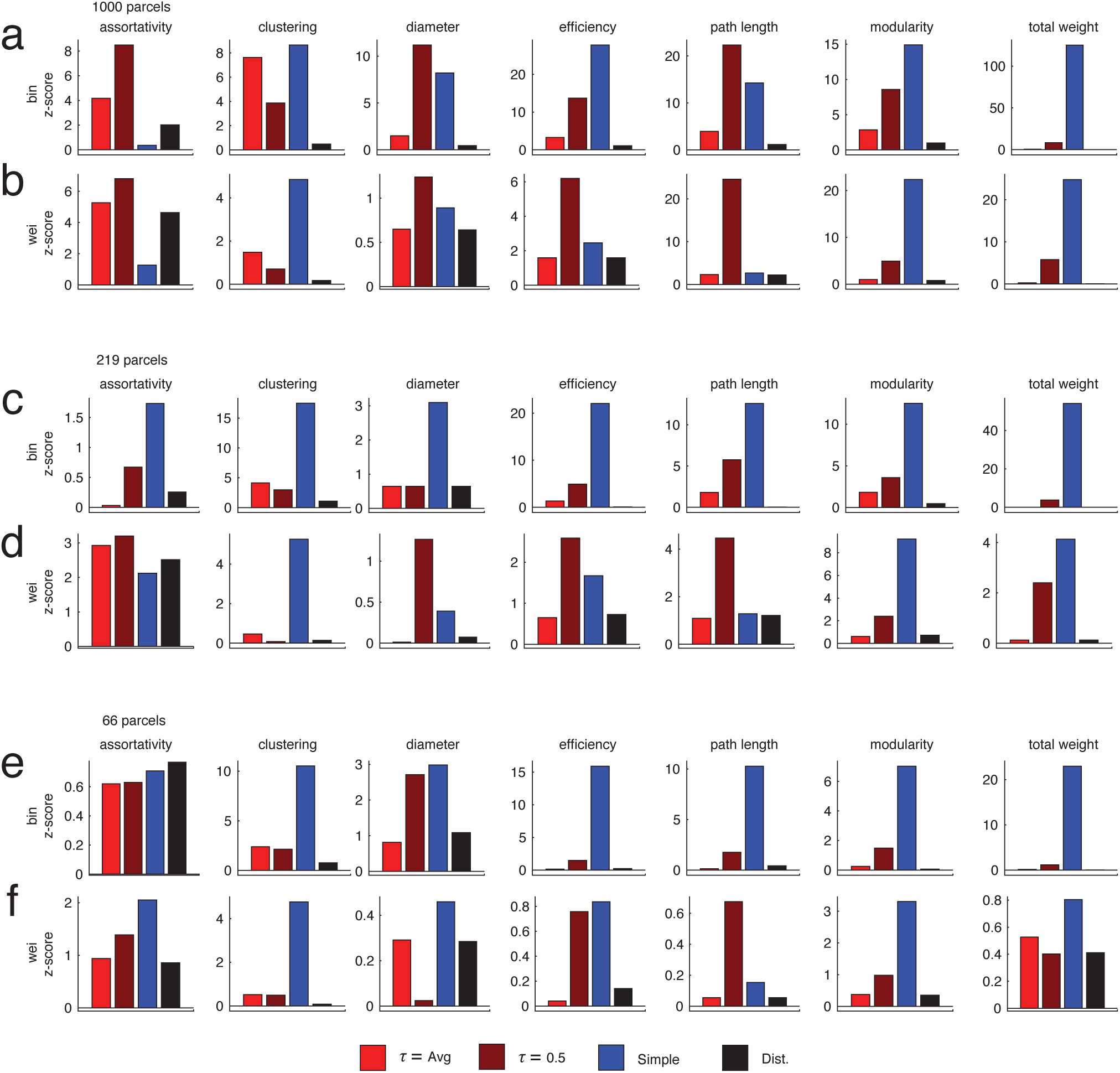
Z-score comparisons of subject and group networks along different network statistics and with different parcellation resolutions. In Fig. S1 we show the ranking of group networks in terms of how well they minimized the difference between themselves and subject-level networks in terms of different network statistics. Those rankings, while informative, could be susceptible to particular biases. Namely, if the different in performance between two models is, small, i.e. effectively the same, the ranking system will still use that negligible difference to rank one of the models higher than the other. Here, we show the raw z-scores used to compute those rankings. In panels *a* and *b*, we show z-scores for weighted and binary networks using the *N* = 1000 node parcellation. Panels *c* + *d* and *e* + *f* show the same for *N* = 219 and *N* = 66 node networks.

